# A single locus underlies variation in *Caenorhabditis elegans* chemotherapeutic responses

**DOI:** 10.1101/2020.03.09.984393

**Authors:** Kathryn S. Evans, Erik C. Andersen

## Abstract

Pleiotropy, the concept that a single gene controls multiple distinct traits, is prevalent in most organisms and has broad implications for medicine and agriculture. Identifying the molecular mechanisms underlying pleiotropy has the power to unveil previously unknown biological connections between seemingly unrelated traits. Additionally, the discovery of pleiotropic genes increases our understanding of both genetic and phenotypic complexity by characterizing novel gene functions. Quantitative trait locus (QTL) mapping has been used to identify several pleiotropic regions in many organisms. However, gene knockout studies are needed to eliminate the possibility of tightly linked, non-pleiotropic loci. Here, we use a panel of 296 recombinant inbred advanced intercross lines of *Caenorhabditis elegans* and a high-throughput fitness assay to identify a single large-effect QTL on the center of chromosome V associated with variation in responses to eight chemotherapeutics. We validate this QTL with near-isogenic lines and pair genome-wide gene expression data with drug response traits to perform mediation analysis, leading to the identification of a pleiotropic candidate gene, *scb-1*. Using deletion strains created by genome editing, we show that *scb-1*, which was previously implicated in response to bleomycin, also underlies responses to other double-strand DNA break-inducing chemotherapeutics. This finding provides new evidence for the role of *scb-1* in the nematode drug response and highlights the power of mediation analysis to identify causal genes.

## INTRODUCTION

Pleiotropy refers to the well established notion that a single gene or genetic variant affects multiple distinct traits (Paaby and Rockman 2013), and the discovery of pleiotropic genes can provide meaningful insights into the molecular mechanisms of these traits (Tyler, Crawford, and Pendergrass 2016). It has become easier to identify pleiotropic genes with the advent of reverse-genetic screens and quantitative trait locus (QTL) mapping (Paaby and Rockman 2013). For example, pleiotropic QTL for diverse growth and fitness traits have been identified in organisms such as yeast (Peltier et al. 2019; Jerison et al. 2017; Cubillos et al. 2011), *Arabidopsis* (Fusari et al. 2017; McKay, Richards, and Mitchell-Olds 2003; El-Assal et al. 2004), *Drosophila* (McGuigan et al. 2014; Brown et al. 2013), and mice (White et al. 2013; Leamy et al. 2014; Lin et al. 2014). These studies have led to important questions in the field of evolutionary genetics regarding the ‘cost of complexity’ (Fisher, n.d.; Orr 2000), as a single mutation might be beneficial for one trait and harmful for another (Wagner and Zhang 2011). Furthermore, human association studies have identified pleiotropic variants associated with different diseases (Sivakumaran et al. 2011; Chesmore, Bartlett, and Williams 2018; Pavlides et al. 2016), highlighting both the ubiquity and importance of certain immune-related genes and oncogenes across unrelated diseases (Gratten and Visscher 2016; Borrello, Degl’Innocenti, and Pierotti 2008). Perhaps the strongest evidence of pleiotropy exists for molecular phenotypes. Large-scale expression QTL (eQTL) mapping studies have identified single regulatory variants that control expression and likely the functions of hundreds of genes at once, opening a window into the mechanisms for how traits are controlled (Rockman, Skrovanek, and Kruglyak 2010; Breitling et al. 2008; Hasin-Brumshtein et al. 2016; Keurentjes et al. 2007; Frank Wolfgang Albert et al. 2018; Frank W. Albert and Kruglyak 2015).

The nematode *Caenorhabditis elegans* provides a tractable metazoan model to identify and study pleiotropic QTL (Paaby and Rockman 2013). A large panel of recombinant inbred advanced intercross lines (RIAILs) derived from two divergent strains, N2 and CB4856 (Rockman and Kruglyak 2009; Andersen et al. 2015), has been leveraged in several linkage mapping analyses (McGrath et al. 2009; Bendesky and Bargmann 2011; Lee et al. 2017; Singh et al. 2016; Zdraljevic et al. 2019; Brady et al. 2019; Zdraljevic et al. 2017; Evans et al. 2018; Andersen et al. 2014; Viñuela et al. 2010; Doroszuk et al. 2009; Snoek et al. 2014; Rodriguez et al. 2012; Glater, Rockman, and Bargmann 2014; Rockman, Skrovanek, and Kruglyak 2010; Zamanian et al. 2018a; Bendesky et al. 2011, 2012; Schmid et al. 2015; Balla et al. 2015; Kammenga et al. 2007; Gutteling, Riksen, et al. 2007; Gutteling, Doroszuk, et al. 2007; Li et al. 2006; Reddy et al. 2009; Seidel et al. 2011; Seidel, Rockman, and Kruglyak 2008). Quantitative genetic analysis using these panels and a high-throughput phenotyping assay (Andersen et al. 2015) has facilitated the discovery of numerous QTL (Zamanian et al. 2018b), several quantitative trait genes (QTG) (Brady et al. 2019) and quantitative trait nucleotides (QTN) (Zdraljevic et al. 2019, 2017) underlying fitness-related traits in the nematode. Additionally, three pleiotropic genomic regions were recently found to influence responses to a diverse group of toxins (Evans et al. 2018). However, overlapping genomic regions might not represent true pleiotropy but could demonstrate the co-existence of tightly linked loci (Paaby and Rockman 2013).

Here, we use linkage mapping to identify a single overlapping QTL on chromosome V that influences the responses to eight chemotherapeutic compounds. We show that these drug-response QTL also overlap with an expression QTL hotspot that contains the gene *scb-1*, previously implicated in bleomycin response (Brady et al. 2019). Although the exact mechanism of *scb-1* is yet unknown, it is hypothesized to act in response to stress (Riedel et al. 2013) and has weak homology to a viral hydrolase (Zhang et al. 2018; Kelley et al. 2015). Together, these data suggest that the importance of *scb-1* expression might extend beyond bleomycin response. We validated the QTL using near-isogenic lines (NILs) and performed mediation analysis to predict that *scb-1* expression explains the observed QTL for five of the eight drugs. Finally, we directly tested the effect of *scb-1* loss of function on chemotherapeutic responses. We discovered that expression of *scb-1* underlies differential responses to several chemotherapeutics that cause double-strand DNA breaks, not just bleomycin. This discovery of pleiotropy helps to further define the role of *scb-1* by expanding its known functions and provides insights into the molecular mechanisms underlying the nematode drug response.

## MATERIALS AND METHODS

### Strains

Animals were grown at 20°C on modified nematode growth media (NGMA) containing 1% agar and 0.7% agarose to prevent burrowing and fed OP50 (Ghosh et al. 2012). The two parental strains, the canonical laboratory strain, N2, and the wild isolate from Hawaii, CB4856, were used to generate all recombinant lines. 208 recombinant inbred advanced intercross lines (RIAILs) generated previously by Rockman *et al.* (Rockman and Kruglyak 2009) (set 1 RIAILs) were phenotyped for expression QTL mapping (detailed below). A second set of 296 RIAILs generated previously by Andersen *et al.* (Andersen et al. 2015) (set 2 RIAILs) was used more extensively for drug phenotyping and linkage mapping. Near-isogenic lines (NILs) were generated by backcrossing a selected RIAIL for several generations (Brady et al. 2019), using PCR amplicons for insertion-deletion (indels) variants to track the introgressed region. NILs were whole-genome sequenced to verify clean introgressions. CRISPR-Cas9-mediated deletions of *scb-1* were described in Brady *et al.* (Brady et al. 2019). All strains are available upon request or from the *C. elegans* Natural Diversity Resource (Cook et al. 2016).

### High-throughput fitness assays for linkage mapping

For dose responses and RIAIL phenotyping, we used a high-throughput fitness assay (HTA) described previously (Andersen et al. 2015). In summary, populations of each strain were passaged and amplified on NGMA plates for four generations. In the fifth generation, gravid adults were bleach-synchronized and 25-50 embryos from each strain were aliquoted into 96-well microtiter plates at a final volume of 50 μL K medium (Boyd, Smith, and Freedman 2012). The following day, arrested L1s were fed HB101 bacterial lysate (Pennsylvania State University Shared Fermentation Facility, State College, PA; (García-González et al. 2017)) at a final concentration of 5 mg/mL in K medium and were grown to the L4 larval stage for 48 hours at 20°C with constant shaking. Three L4 larvae were sorted into new 96-well microtiter plates containing 10 mg/mL HB101 bacterial lysate, 50 μM kanamycin, and either diluent (1% water or 1% DMSO) or drug dissolved in the diluent using a large-particle flow cytometer (COPAS BIOSORT, Union Biometrica; Holliston, MA). Sorted animals were grown for 96 hours at 20°C with constant shaking. The next generation of animals and the parents were treated with sodium azide (50 mM in 1X M9) to straighten their bodies for more accurate length measurements. Animal length (median.TOF), optical density (median.norm.EXT), and brood size (norm.n) were quantified for each well using the COPAS BIOSORT. Phenotypic measurements collected by the BIOSORT were processed and analyzed using the R package *easysorter* (Shimko and Andersen 2014) as described previously (Brady et al. 2019).

### Dose-response assays

Four genetically divergent strains (N2, CB4856, JU258, and DL238) were treated with increasing concentrations of each of the eight drugs using the HTA described above. The dose of each drug that provided a reproducible drug-specific effect that maximizes between-strain variation while minimizing within-strain variation across the three traits was selected for the linkage mapping experiments. The chosen concentrations are as follows: 100 μM amsacrine hydrochloride (Fisher Scientific, #A277720MG) in DMSO, 50 μM bleomycin sulfate (Fisher, #50-148-546) in water, 2 μM bortezomib (VWR, #AAJ60378-MA) in DMSO, 250 μM carmustine (Sigma, #1096724-75MG) in DMSO, 500 μM cisplatin (Sigma, #479306-1G) in K media, 500 μM etoposide (Sigma, #E1383) in DMSO, 500 μM puromycin dihydrochloride (VWR, #62111-170) in water, and 150 μM silver nitrate (Sigma-Aldrich, #S6506-5G) in water.

### Linkage mapping

Set 1 and set 2 RIAILs were phenotyped in each of the eight drugs and controls using the HTA described above. Linkage mapping was performed on each of the drug and gene expression traits using the R package *linkagemapping* (https://github.com/AndersenLab/linkagemapping) as described previously (Brady et al. 2019). The cross object derived from the whole-genome sequencing of the RIAILs containing 13,003 SNPs was loaded using the function *load_cross_obj(“N2xCB4856cross_full”)*. The RIAIL phenotypes were merged into the cross object using the *merge_pheno* function with the argument *set* = 1 for expression QTL mapping and *set* = 2 for drug phenotype mapping. A forward search (*fsearch* function) adapted from the *R/qtl* package (Broman et al. 2003) was used to calculate the logarithm of the odds (LOD) scores for each genetic marker and each trait as *-n(ln(1-R^2^)/2ln(10))* where R is the Pearson correlation coefficient between the RIAIL genotypes at the marker and trait phenotypes (Bloom et al. 2013). A 5% genome-wide error rate was calculated by permuting the RIAIL phenotypes 1000 times. The marker with the highest LOD score above the significance threshold was selected as the QTL then integrated into the model as a cofactor and mapping was repeated iteratively until no further significant QTL were identified. Finally, the *annotate_lods* function was used to calculate the effect size of each QTL and determine 95% confidence intervals defined by a 1.5 LOD drop from the peak marker using the argument *cutoff* = *proximal*.

### Modified HTA for NIL validation

NILs and *scb-1* deletion strains were tested using a modified version of the HTA detailed above. Strains were propagated for two generations, bleach-synchronized in three independent replicates, and titered at a concentration of 25-50 embryos per well of a 96-well microtiter plate. The following day, arrested L1s were fed HB101 bacterial lysate at a final concentration of 5 mg/mL with either diluent or drug. After 48 hours of growth at 20°C with constant shaking, nematodes were treated with sodium azide (5 mM in water) prior to analysis of animal length and optical density using the COPAS BIOSORT. As only one generation of growth is observed, brood size was not calculated. A single trait (median.EXT) was chosen to represent animal growth generally, as the trait is defined by optical density over length. Because of the modified timing of the drug delivery, lower drug concentrations were needed to see the previous effect. The selected doses are as follows: 12.5 μM amsacrine in DMSO, 12.5 μM bleomycin in water, 2 μM bortezomib in DMSO, 250 μM carmustine in DMSO, 125 μM cisplatin in K media, 62.5 μM etoposide in DMSO, 300 μM puromycin in water, and 100 μM silver in water.

### Expression QTL analysis

Microarray data for gene expression using 15,888 probes were previously collected from synchronized young adult populations of 209 set 1 RIAILs (Rockman, Skrovanek, and Kruglyak 2010). Expression data were corrected for dye effects and probes with variants were removed (Andersen et al. 2014). Linkage mapping was performed as described above for the remaining 14,107 probes, and a significance threshold was determined using a permutation-based False Discovery Rate (FDR). FDR was calculated as the ratio of the average number of genes across 10 permutations expected by chance to show a maximum LOD score greater than a particular threshold vs. the number of genes observed in the real data with a maximum LOD score greater than that threshold. We calculated the FDR for a range of thresholds from 2 to 10, with increasing steps of 0.01, and set the threshold so that the calculated FDR was less than 5%.

Local eQTL were defined as linkages whose peak LOD scores were within 1 Mb of the starting position of the probe (Rockman, Skrovanek, and Kruglyak 2010). eQTL hotspots were identified by dividing the genome into 5 cM bins and counting the number of distant eQTL that mapped to each bin. Significance was determined as bins with more eQTL than the Bonferroni-corrected 99^th^ percentile of a Poisson distribution with a mean of 3.91 QTL (total QTL / total bins) (Brem et al. 2002; Evans et al. 2018; Rockman, Skrovanek, and Kruglyak 2010).

### Mediation analysis

A total of 159 set 1 RIAILs were phenotyped in each of the eight drugs and controls using the HTA described above. Mediation scores were calculated with bootstrapping using the *mediate* function from the *mediation* R package (version 4.4.7) (Tingley et al. 2014) for each QTL identified from the set 2 RIAILs and all 49 probes (including *scb-1,* A_12_P104350) within the chromosome V eQTL hotspot using the following models:

1. Mediator model: *lm(expression ~ genotype)*
2. Outcome model: *lm(phenotype ~ expression + genotype)*

The output of the *mediate* function can be summarized as follows: the total effect of genotype on phenotype, ignoring expression (*tau.coef*); the direct effect of genotype on phenotype, while holding expression constant (*z0*); the estimated effect of expression on phenotype (*d0*); the proportion of the total effect that can be explained by expression data (*n0*) The final mediation score is determined as the proportion of the total QTL effect that can be attributed to gene expression (*n0*). The likelihood of *scb-1* mediating a given QTL effect was calculated relative to the other 74 probes in the region. Traits in which *scb-1* was at or above the 95^th^ percentile of this distribution were prioritized over other traits.

### Statistical analysis

Broad-sense heritability was calculated from the dose response phenotypes using the *lmer* function in the *lme4* R package (Bates et al. 2014) with the formula *phenotype* ~ *1* + (*1*|*strain*) for each dose. All statistical tests of phenotypic differences between strains were performed using the *TukeyHSD* function (R Core Team 2017) on an ANOVA model with the formula *phenotype* ~ *strain*.

### Data Availability

**File S1** contains the results of the original dose response HTA. **File S2** contains the residual phenotypic values for all 159 set 1 RIAILs, 296 set 2 RIAILs, and parent strains (N2 and CB4856) in response to all eight chemotherapeutics. **File S3** contains the linkage mapping results for the set 2 RIAILs for all 24 drug-response traits tested in the HTA. **File S4** contains the genotype of the NILs in the study. **File S5** contains the raw pruned phenotypes for the NIL dose response with the modified HTA. **File S6** contains the pairwise statistical significance for all strains and high-throughput assays. **File S7** contains the microarray expression data for 14,107 probes from Rockman *et al.* 2010. **File S8** contains the linkage mapping results for the expression data obtained with the set 1 RIAILs. **File S9** contains the location of each eQTL hotspot and a list of genes with an eQTL in each hotspot. **File S10** contains the linkage mapping results from the set 1 RIAILs for all 24 drug-response traits tested in the HTA. **File S11** contains the pairwise mediation estimates for all QTL and all 75 probes. **File S12** contains the raw pruned phenotypes for the *scb-1* deletion modified HTA. The datasets and code for generating figures can be found at https://github.com/AndersenLab/scb1_mediation_manuscript. Supplemental material available at Figshare.

## RESULTS

### Natural variation on chromosome V underlies differences in responses to several chemotherapeutics

We measured *C. elegans* development and chemotherapeutic sensitivity as a function of animal length (TOF), optical density (EXT), and brood size (n) with a high-throughput assay developed using the COPAS BIOSORT (see Methods) (Zdraljevic et al. 2019; Brady et al. 2019; Evans et al. 2018; Zdraljevic et al. 2017; Andersen et al. 2015). We exposed four genetically divergent strains (N2, CB4856, JU258, and DL238) to increasing doses of eight chemotherapeutic compounds. Five of these compounds (bleomycin, carmustine, etoposide, amsacrine, and cisplatin) are known to cause double-strand DNA breaks and/or inhibit DNA synthesis (Dorr 1992; Dasari and Tchounwou 2014; Montecucco, Zanetta, and Biamonti 2015; Ketron et al. 2012; Nikolova et al. 2017). The remaining three compounds either inhibit protein synthesis (puromycin) (Azzam and Algranati 1973), inhibit the proteosome and subsequent protein degradation (bortezomib) (Piperdi et al. 2011), or cause cellular toxicity in a poorly defined way (silver nitrate) (Kaplan, Akalin Ciftci, and Kutlu 2016) (**Table 1**). In the presence of each drug, nematodes were generally shorter, less optically dense, and produced smaller broods compared to non-treated nematodes (**Figure S1, File S1**). We observed significant phenotypic variation among strains and identified a substantial heritable genetic component for most traits (average *H*^2^ = 0.51 +/− 0.24).

**Table 1:** Main mechanism of action for eight chemotherapeutic drugs.

| **Drug** | **Drug class** | **Mechanism of action** |
| --- | --- | --- |
| Amsacrine | Topoisomerase inhibitors | DNA intercalation and inhibition of topoisomerase II, causing DNA double-strand breaks, cell cycle arrest, and cell death |
| Bleomycin | Antitumor antibiotic | Forms complexes with iron that reduce molecular oxygen to form free radicals which in turn cause DNA single- and double-strand breaks |
| Bortezomib | Proteasome inhibitors | Reversibly inhibits the 26S proteasome and inhibits nuclear factor (NF)-kappaB resulting in disruption of various cell signaling pathways, cell cycle arrest, and cell death. |
| Carmustine | Alkylating agents | Alkylates and cross-links DNA causing cell cycle arrest and cell death |
| Cisplatin | Alkylating agents | Alkylates and cross-links DNA causing cell cycle arrest and cell death |
| Etoposide | Topoisomerase inhibitors | Binds to and inhibits topoisomerase II causing an increase of DNA single- and double-strand breaks, cell cycle arrest, and cell death |
| Puromycin | Aminonucleoside antibiotic | Acts as analog of 3' terminal end of aminoacyl-tRNA and incorporates itself into growing polypeptide chain causing premature termination and inhibition of protein synthesis |
| Silver | NA | Multi-faceted induction of apoptosis |

We exposed a panel of 296 RIAILs (set 2 RIAILs, see Methods) to all eight chemotherapeutics at a selected concentration that both maximizes among-strain and minimizes within-strain phenotypic variation (**File S2**). Linkage mapping for all three traits for each of the eight drugs (total of 24 traits) identified 56 QTL from 23 traits (one trait had no significant QTL), several of which have been identified previously (Brady et al. 2019; Evans et al. 2018; Zdraljevic et al. 2017) (**File S3, Figure S2**). Strikingly, a QTL on the center of chromosome V was linked to variation in responses to all eight compounds (**Figure 1**). In all cases, the CB4856 allele on chromosome V is associated with greater resistance to the drug than the N2 allele (**Figure S2, File S2, File S3**). We previously identified this genomic interval as a QTL hotspot, defined as a region heavily enriched for toxin-response QTL (Evans et al. 2018). Because several of the chemotherapeutics share a similar mechanism of action, a single pleiotropic gene might underlie the observed QTL for multiple drugs.

**Figure 1.**
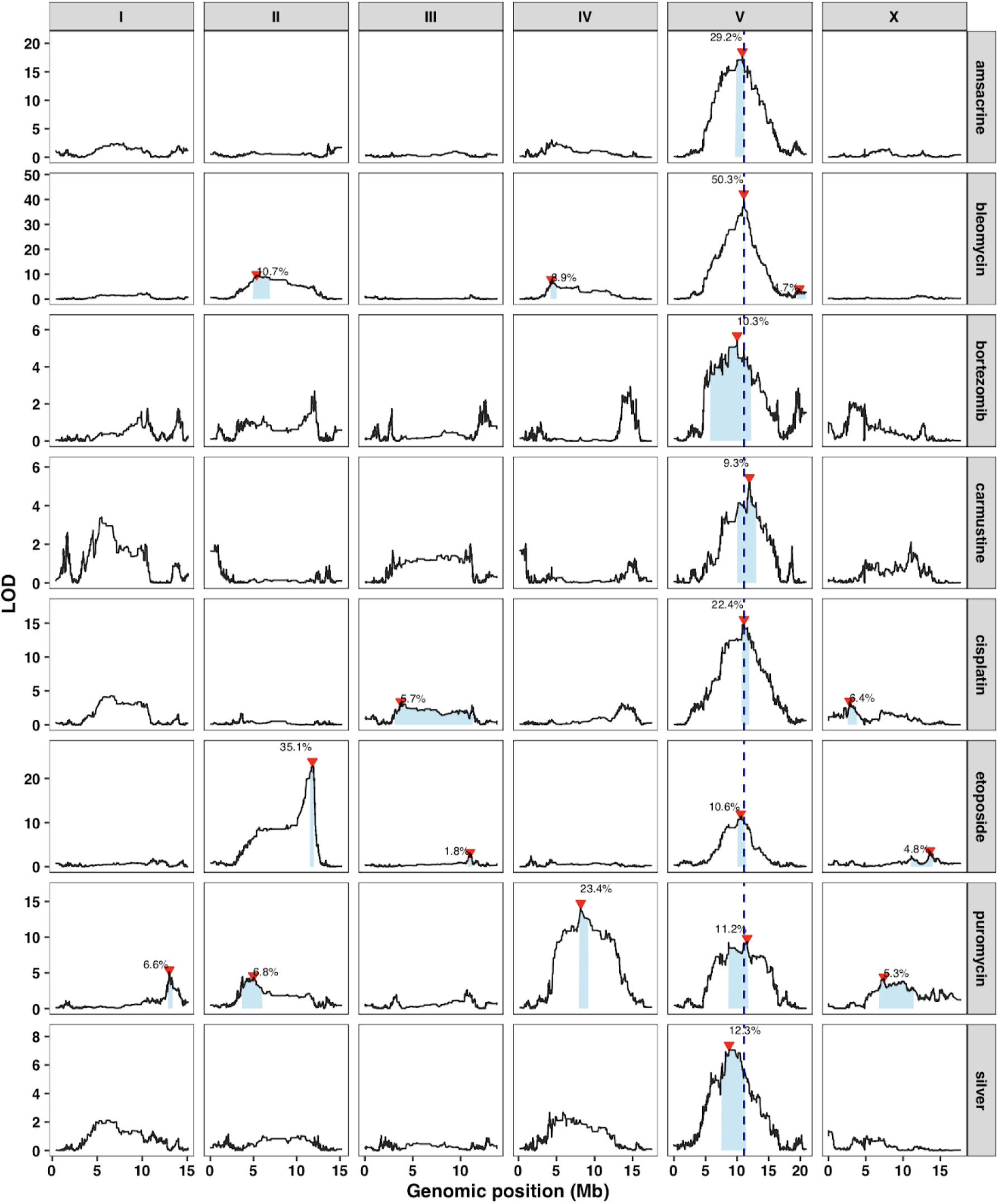
A large-effect QTL on the center of chromosome V underlies responses to several chemotherapeutics. Linkage mapping results for a representative trait for each drug are shown. Genomic position (x-axis) is plotted against the logarithm of the odds (LOD) score (y-axis) for 13,003 genomic markers. Each significant QTL is indicated by a red triangle at the peak marker, and a blue rectangle shows the 95% confidence interval around the peak marker. The percentage of the total variance in the RIAIL population that can be explained by each QTL is shown above the QTL. The dotted vertical line represents the genomic position of *scb-1*.

In order to isolate and validate the effect of this QTL, we constructed reciprocal near-isogenic lines (NILs) by introgressing a genomic region on chromosome V from the resistant CB4856 strain into the sensitive N2 background and vice versa (**File S4**). We used a modified high-throughput assay (see Methods) to measure length and optical density of a population of animals grown in the presence of the drug for 48 hours (from larval stages L1 to L4). In this modified assay, less drug was required to observe the same phenotypic effect as before and a single trait (median.EXT) that combines both optical density and length serves as a proxy for the complexity of animal growth (**Figure S3, File S5**). For all eight chemotherapeutics tested, the strain with the N2 introgression was significantly more sensitive than its CB4856 parent and/or the strain with the CB4856 introgression was significantly more resistant than its N2 parent (**Figure 2, File S5, File S6**). These data confirm that one or more genetic variant(s) within this region on chromosome V cause increased drug sensitivities in N2.

**Figure 2.**
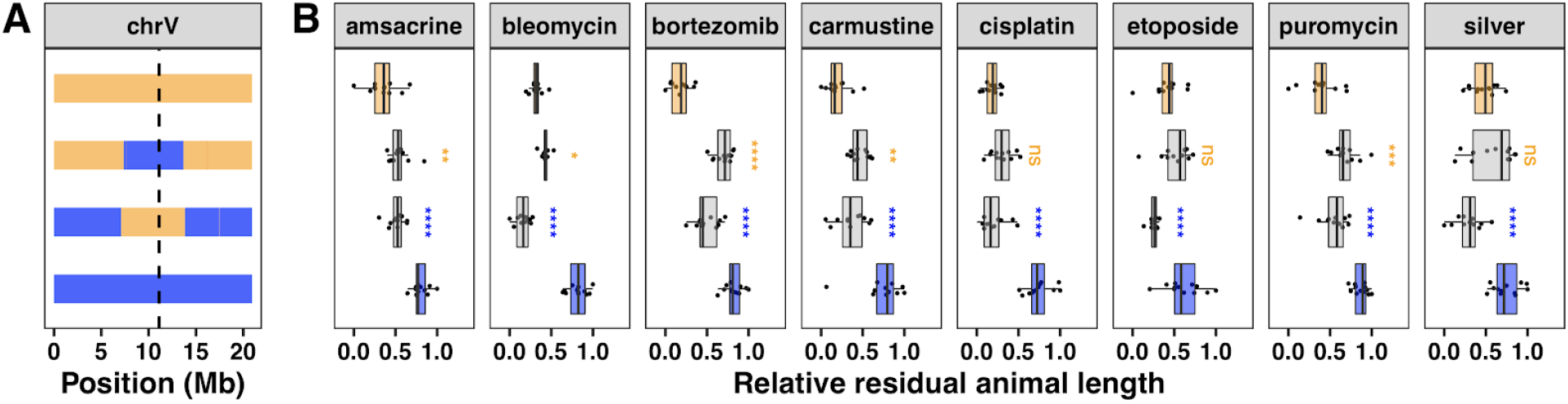
Near-isogenic lines validate the chromosome V QTL. (A) NIL genotypes on chromosome V are shown, colored orange (N2) and blue (CB4856). From top to bottom, strains are N2, ECA232, ECA1114, and CB4856. The dotted vertical line represents the location of *scb-1*. (B) NIL phenotypes are plotted as Tukey box plots with strain (y-axis) by relative residual animal length (x-axis). Statistical significance of each NIL compared to its parental strain (ECA232 to N2 and ECA1114 to CB4856) is shown above each NIL and colored by the parent strain against which it was tested (ns = non-significant (p-value > 0.05); *, **, ***, and *** = significant (p-value < 0.05, 0.01, 0.001, or 0.0001, respectively).

### Expression QTL mapping identifies a hotspot on the center of chromosome V

Genetic variation can affect a phenotype most commonly through either modifications of the amino acid sequence that lead to altered protein function (or even loss of function) or changes in the expression level of the protein. In the latter case, measuring the intermediate phenotype (gene expression) can be useful in elucidating the mechanism by which genetic variation causes phenotypic variation. More specifically, cases with overlap between expression QTL (eQTL) and drug-response QTL suggest that a common variant could underlie both traits and provide evidence in support of causality for the candidate gene in question (Sasaki, Frommlet, and Nordborg 2018; Huang et al. 2015).

To identify such cases of overlap between expression QTL and the drug-response QTL on chromosome V, we need genome-wide expression data for the RIAILs. Expression of 15,888 probes were previously measured using microarrays for a panel of 208 RIAILs (set 1 RIAILs, see Methods) between N2 and CB4856 (Rockman and Kruglyak 2009) (**File S7**). The variation in gene expression was used as a phenotypic trait to identify eQTL using linkage mapping with 1,455 variants (Rockman, Skrovanek, and Kruglyak 2010). Rockman *et al.* identified 2,309 eQTL and three regions with significantly clustered distant eQTL (eQTL hotspots), suggesting that these regions are pleiotropic, wherein one or more variant(s) are affecting expression of multiple genes. We performed whole-genome sequencing for these strains and identified 13,003 informative variants (Brady et al. 2019). Linkage mapping with these variants for the 14,107 probes without genetic variation in CB4856 (Andersen et al. 2014) identified 2,540 eQTL associated with variation in expression of 2,196 genes (**Figure 3A, File S8**). These eQTL have relatively large effect sizes compared to the drug-response QTL. On average, each eQTL explains 23% of the phenotypic variance in gene expression among the RIAIL population. Half of the eQTL (50.2%; 1,276) mapped within 1 Mb of the gene whose expression was measured and were classified as local (see Methods). The other half (49.7%; 1,264) were found distant from their respective gene, and over a third (37%; 940) were found on different chromosomes entirely. In general, eQTL effect sizes increased, max LOD scores decreased, and confidence intervals become smaller compared to the original mapping results (**File S8**). These differences and the additional eQTL observed between this analysis and the original are likely due to the integration of new genetic markers.

**Figure 3.**
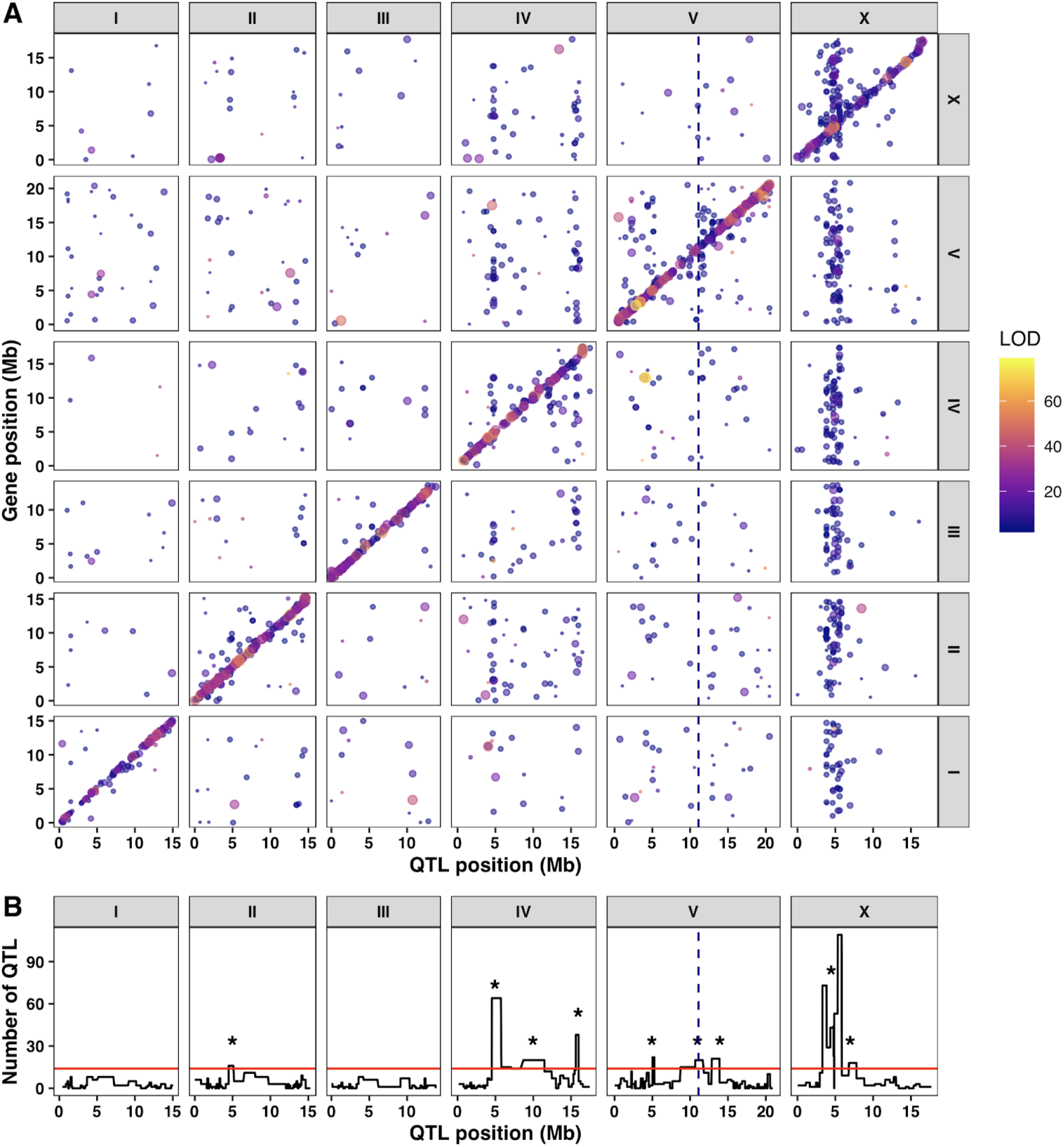
Expression QTL mapping identifies several hotspots. (A) The genomic locations of the eQTL peaks (x-axis) are plotted against the genomic locations of the probe (y-axis). The size of the point corresponds to the effect size of the QTL. eQTL are colored by the LOD score, increasing from purple to pink to yellow. The diagonal band represents local eQTL, and vertical bands represent eQTL hotspots. (B) Quantification of eQTL hotspots identified by overlapping distant eQTL. The number of distant eQTL (y-axis) in each 5 cM bin across the genome (x-axis) is shown. Bins above the red line are significant and marked with an asterisk. The dotted vertical line represents the genomic position of *scb-1*.

In total, we identified nine eQTL hotspots (**Figure 3B, File S9**). Three of which were previously identified on chromosome IV and X (Rockman, Skrovanek, and Kruglyak 2010). Notably, three of the eQTL hotspots overlap with the previously identified drug-response QTL hotspots on chromosomes IV and V (Evans et al. 2018). The overlap of these eQTL and drug-response QTL hotspots could provide strong candidate genes whose expression underlies the differences in nematode drug responses generally. Expression of one gene of interest, *scb-1*, has been previously implicated in response to bleomycin (Brady et al. 2019) and resides within the eQTL hotspot region on chromosome V (**File S9**). Together with the putative role for *scb-1* as a hydrolase (Zhang et al. 2018; Kelley et al. 2015; Brady et al. 2019), these data suggest that variation in expression of *scb-1* and responses to these eight chemotherapeutics (including bleomycin) could be mechanistically linked.

### Mediation analysis suggests *scb-1* expression plays a role in responses to several chemotherapeutics

Mediation analysis seeks to explain the relationship between an independent and dependent variable by including a third (non-observable) intermediary variable. We can use mediation analysis to understand how certain genetic variants on chromosome V (independent variable) affect drug responses (dependent variable) through differential gene expression of genes within the eQTL hotspot (mediator variable) (**Figure S4**). We measured brood size, animal length, and optical density in response to all eight chemotherapeutics in the set 1 RIAILs and performed linkage mapping for these traits (**File S2, File S10, Figure S5**). Although the power to detect QTL with these strains is lower than in our original mapping set (set 2 RIAILs) (Andersen et al. 2015), we still identified overlapping QTL on chromosome V for half of the drugs tested (bleomycin, cisplatin, silver, and amsacrine) (**Figure S5**). We also detected putative small-effect QTL just below the permutation-based threshold for at least one other chemotherapeutic (**File S10**).

We calculated the proportion of QTL effects that can be explained by variation in expression of *scb-1* compared to the other 48 genes in the chromosome V eQTL hotspot using mediation analysis (see Methods). We estimated that expression of *scb-1* mediates 61.3% of the chromosome V QTL for bleomycin response (**Figure 4, File S11**). Moreover, out of all 49 genes in the region, *scb-1* was a clear mediation score outlier, falling in the 99^th^ percentile of all local genes. Of the remaining seven chemotherapeutics, we showed that amsacrine, cisplatin, etoposide, and puromycin all showed moderate evidence of *scb-1* mediation, with *scb-1* falling near or above the 95^th^ percentile of mediation estimates for all local genes (**Figure 4, File S11**). In fact, *scb-1* was the only gene with a significant mediation estimate within the drug-response QTL confidence interval for cisplatin. The opposite result was seen for bortezomib, carmustine, and silver, in which all showed strong evidence against *scb-1* mediation (**Figure 4, File S11**). This *in silico* approach indicated that expression of *scb-1* might be an intermediate link between genetic variation on chromosome V and responses to five of the eight chemotherapeutic drugs tested.

**Figure 4.**
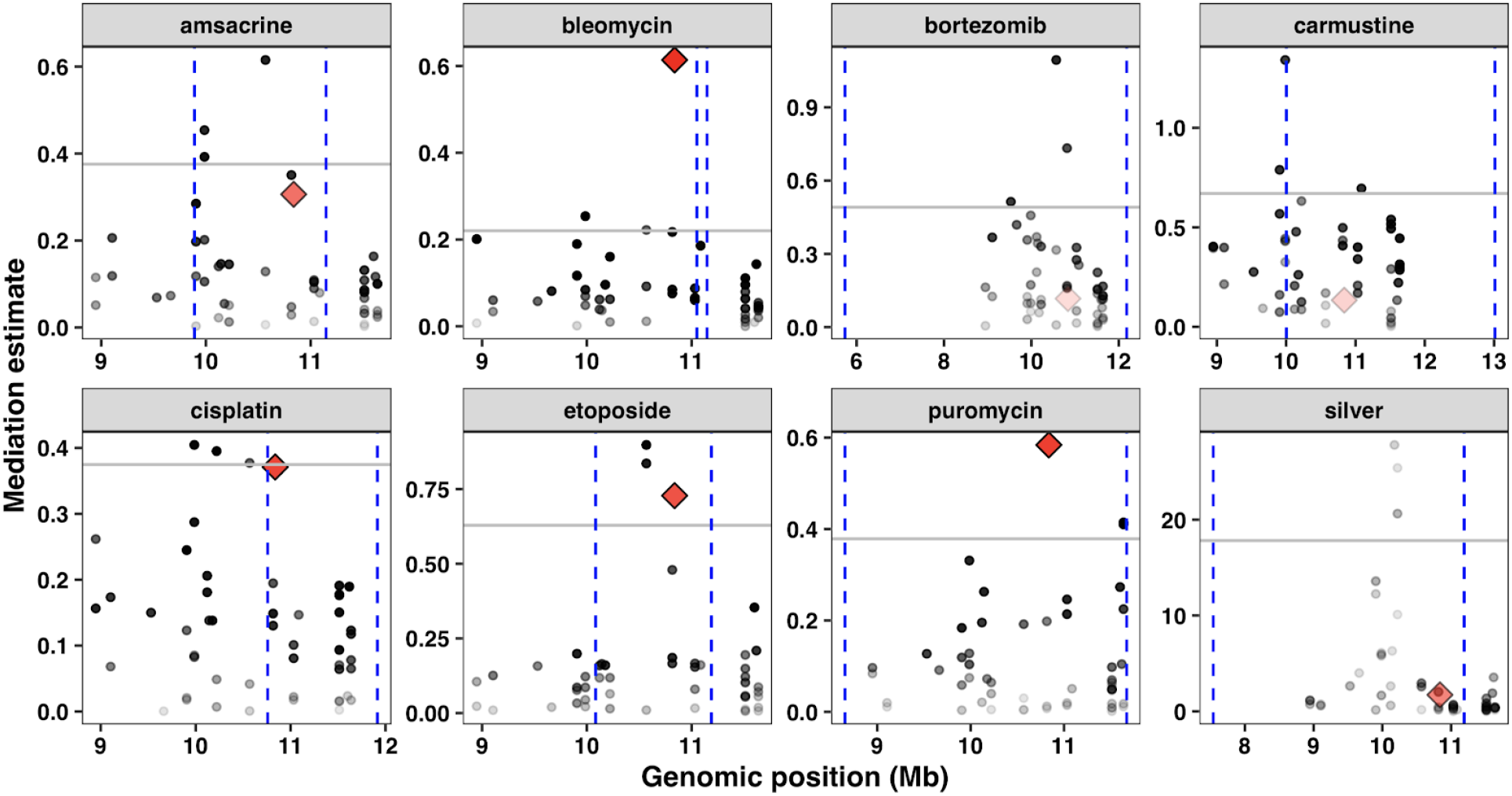
Local mediation analysis for *scb-1*. Mediation estimates calculated as the fraction of the total QTL effect explained by differences in expression of each gene (y-axis) are plotted against genomic position of the gene (x-axis) on chromosome V for 74 probes surrounding *scb-1* (red diamond). The 95th percentile of the distribution of mediation estimates for each trait are represented by the horizontal grey lines. The confidence intervals for QTL are shown with the vertical blue dotted lines. The confidence of the estimate increases (p-value decreases) as points become less transparent.

### Expression of *scb-1* mediates responses to several chemotherapeutics that cause double-strand DNA breaks

To empirically test whether *scb-1* expression modulates the chromosome V QTL effect for each drug, we exposed two independently derived strains with *scb-1* deletions (Brady et al. 2019) to each chemotherapeutic (**Figure 5, Figure S6, File S6, File S12**). Because RIAILs with the CB4856 allele on chromosome V express higher levels of *scb-1* than RIAILs with the N2 allele (**File S7, File S8**), we expect that loss of *scb-1* will cause increased drug sensitivity in the CB4856 background but might not have an effect in the N2 background. We validated the results of Brady *et al.* and confirmed that ablated *scb-1* expression causes robust sensitivity to bleomycin in both N2 and CB4856 (**Figure 5, Figure S6, File S6, File S12**). We also observed similarly increased sensitivity to cisplatin with *scb-1* deletions in both backgrounds. Furthermore, removing *scb-1* shows moderately increased sensitivity in the CB4856 background for amsacrine and in the N2 background for carmustine. The remaining four drugs did not show a significantly different phenotype between the parental N2 and CB4856 strains, suggesting these traits might be less reproducible or that *scb-1* variation does not underlie these drug differences. Overall, these results provide evidence for the pleiotropic effect of *scb-1*, which appears to mediate responses to four different drugs.

**Figure 5.**
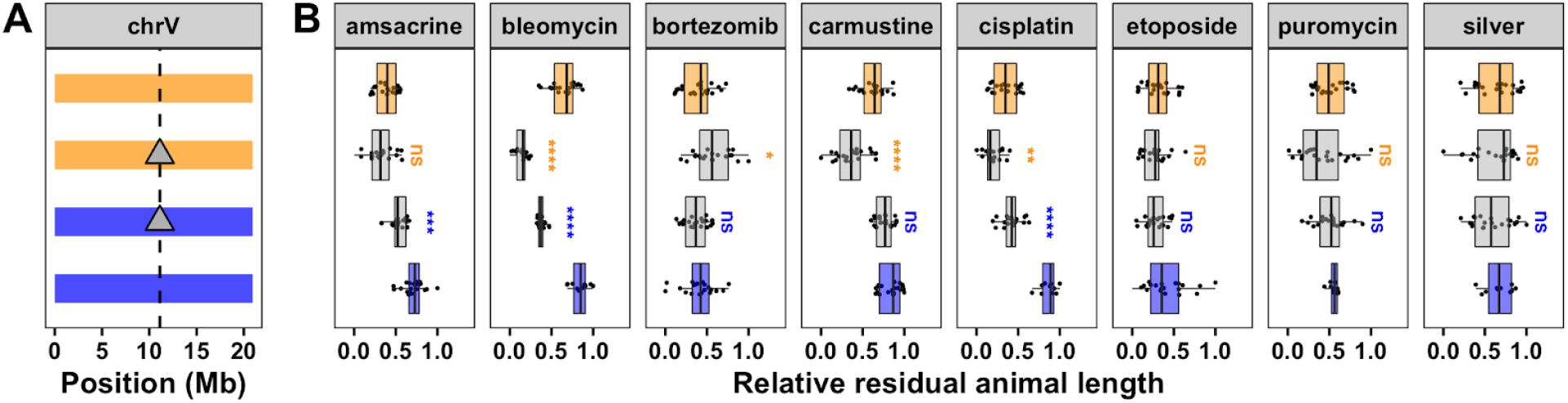
Testing the role of *scb-1* in drug responses. (A) Strain genotypes on chromosome V are shown, colored orange (N2) and blue (CB4856). From top to bottom, strains are N2, ECA1132, ECA1134, and CB4856. Deletion of *scb-1* is indicated by a grey triangle. The dotted vertical line represents the location of *scb-1*. (B) Phenotypes are plotted as Tukey box plots with strain (y-axis) by relative residual animal length (x-axis). Statistical significance of each deletion strain compared to its parental strain (ECA1132 to N2 and ECA1134 to CB4856) is shown above each strain and colored by the parent strain against which it was tested (ns = non-significant (p-value > 0.05); *, **, ***, and *** = significant (p-value < 0.05, 0.01, 0.001, or 0.0001, respectively).

## DISCUSSION

In this study, we identified overlapping QTL on the center of chromosome V that influence sensitivities to eight chemotherapeutic drugs. Because five of these drugs are known to cause double-strand DNA breaks, we hypothesized that this genomic region might be pleiotropic – a single shared genetic variant affects the responses to each drug. Because this variant might affect drug responses by regulating gene expression levels, we looked for the co-existence of drug-response QTL and expression QTL on chromosome V. We identified 2,540 eQTL and nine eQTL hotspots, including a region on the center of chromosome V. We calculated the mediation effect of all 49 genes with an eQTL that maps to this hotspot region and identified *scb-1* as a candidate gene whose expression influences the responses to several chemotherapeutics. We used CRISPR-Cas9-mediated *scb-1* deletion strains to empirically validate the role of *scb-1* in the chemotherapeutic response. In addition to bleomycin (Brady et al. 2019), we discovered that responses to cisplatin, amsacrine, and carmustine are mediated by *scb-1* expression. In this study, we found evidence that overlapping QTL are representative of pleiotropy at the gene level and further elucidated the function of *scb-1* as a potential response to double-strand DNA break stress.

### Mediation of drug-response QTL using gene expression to identify causal genes

Mediation analysis often suggests potential candidate genes that underlie different traits (Huang et al. 2015; Sasaki, Frommlet, and Nordborg 2018) and could be applied to drug responses. Using *C. elegans* strains and high-throughput assays, we can rapidly validate hypotheses generated by mediation analysis. Three of the eight chemotherapeutics that map to an overlapping drug-response QTL and were potentially mediated by *scb-1* were validated using targeted deletion strains.

Although mediation analysis provided moderate evidence that expression of *scb-1* may also play a role in sensitivity to etoposide and puromycin, we observed no experimental evidence of this relationship. Additionally, we have evidence that expression of *scb-1* might mediate response to carmustine. However, mediation analysis disagrees. The discrepancy between the mediation analysis and validation of causality using targeted deletion strains could be partially explained by one of several possibilities. First, different traits were measured in each experiment. The mediation analysis used traits measured over 96 hours of growth in drug conditions spanning two generations, but the causality test used traits measured over 48 hours of growth in drug conditions within one generation. Second, the precision of our mediation estimates was likely reduced by the poor quality drug traits for the set 1 RIAIL panel (Andersen et al. 2015). Indeed, bortezomib, carmustine, etoposide, and puromycin did not map to the center of chromosome V using the set 1 RIAILs (**Figure S5**). Expression data for the set 2 RIAIL panel would likely generate more accurate mediation estimates, especially if data were collected using RNA sequencing to avoid the inherent reference bias of microarray data (Zhao et al. 2014). Third, our mediation analysis was performed using expression data collected in control conditions and phenotype data collected in drug conditions. This analysis will only provide evidence of mediation if the baseline expression differences affect an individual’s response to the drug. Collecting expression data from drug-treated nematodes could help us learn more about how gene expression varies in response to treatment with the chemotherapeutic. Finally, as we only directly assessed the complete loss of *scb-1* in drug sensitivity, it is still possible that reduction of function (or change in function) due to a single nucleotide variant or other structural variation in CB4856 could validate the role of *scb-1* in responses to these drugs.

This study demonstrates the power of pairing genome-wide gene expression data with QTL mappings for other traits using simple colocalization as well as more complex mediation analysis techniques. In addition to providing a resource for candidate gene prioritization within a QTL interval, mediation analysis can help to identify the mechanism by which genetic variation causes phenotypic differences. This type of approach could be even more powerful using genome-wide association (GWA) where the decreased linkage disequilibrium between variants generates smaller genomic confidence intervals. Smaller intervals have fewer spurious overlapping eQTL, which could help to narrow the list of candidate genes. Although mediation analysis is only effective if a change in expression is observed and might not be useful for identifying effects from protein-coding variation, many current studies show that the majority of genetic variants associated with complex traits lie in regulatory regions (Hindorff et al. 2009). Whole-genome expression analysis could provide the missing link to the identification of causal genes underlying complex traits.

### New evidence for the pleiotropic function of *scb-1*

We identified eight chemotherapeutics with a QTL that mapped to a genomic region defined as a QTL hotspot on the center of chromosome V (Evans et al. 2018). Multiple genes in close proximity, each regulating an aspect of cellular growth and fitness, might underlie each QTL independently. Alternatively, genetic variation within a single gene might regulate responses to multiple (or all) of the eight drugs tested if the mechanisms of action were shared (*e.g.* repair of double-strand DNA breaks). Expression of *scb-1*, a gene previously implicated in modulating the nematode’s response to bleomycin, was found to reduce sensitivity to half of the drugs tested. This pleiotropic effect of *scb-1* provides new evidence for the function of the gene and possible molecular mechanisms underlying nematode drug responses. It is hypothesized that SCB-1 might function as a hydrolase that breaks down compounds like bleomycin (Brady et al. 2019) or plays a role in the nematode stress response (Riedel et al. 2013). Both hypotheses are consistent with our data, explaining why nematodes with low expression of *scb-1* are highly sensitive to the compound. Furthermore, all four of these chemotherapeutics, whose responses are mediated by expression of *scb-1*, are known to cause double-strand DNA breaks (Dorr 1992; Dasari and Tchounwou 2014; Ketron et al. 2012; Nikolova et al. 2017). Although the results for bortezomib, puromycin, and silver were inconclusive, we found no clear evidence that expression of *scb-1* dictates their responses. Together, these data suggest a potential role for *scb-1* specifically in response to stress induced by double-strand DNA breaks. However, the lack of sensitivity in etoposide, which also causes double-strand DNA breaks (Montecucco, Zanetta, and Biamonti 2015), indicates that this response might be more complex.

The exact variant that causes the differential expression of *scb-1* is still unknown. Importantly, *scb-1* lies within an eQTL hotspot region where it is hypothesized that genetic variation at a single locus might regulate expression of the 49 nearby genes. It is possible that the same causal variant that regulates expression of *scb-1* could also underlie the QTL for the remaining four chemotherapeutics through differential expression of other nearby essential genes. For example, mediation analysis for both bortezomib and etoposide indicated that expression variation of a dehydrogenase (*D1054.8*) may underlie their differential responses (File S11). Overall, our study highlights the power of using mediation analysis to connect gene expression to organismal traits and describes a novel function for the pleiotropic gene *scb-1*.

## Acknowledgments

We would like to thank members of the Andersen Lab for helpful comments on the manuscript. This work was supported by an American Cancer Society Research Scholar Grant (127313-RSG-15-135-01-DD) to E.C.A. Additionally, K.S.E. received support from the NSF-Simons Center for Quantitative Biology at Northwestern University (awards Simons Foundation/SFARI 597491-RWC and the National Science Foundation 1764421).

## References

Albert, Frank W., and Leonid Kruglyak. 2015. “The Role of Regulatory Variation in Complex Traits and Disease.” Nature Reviews. Genetics 16 (4): 197–212.

Albert, Frank Wolfgang, Joshua S. Bloom, Jake Siegel, Laura Day, and Leonid Kruglyak. 2018. “Genetics of Trans-Regulatory Variation in Gene Expression.” eLife 7 (July). https://doi.org/10.7554/eLife.35471.

Andersen, Erik C., Joshua S. Bloom, Justin P. Gerke, and Leonid Kruglyak. 2014. “A Variant in the Neuropeptide Receptor Npr-1 Is a Major Determinant of Caenorhabditis Elegans Growth and Physiology.” PLoS Genetics 10 (2): e1004156.

Andersen, Erik C., Tyler C. Shimko, Jonathan R. Crissman, Rajarshi Ghosh, Joshua S. Bloom, Hannah S. Seidel, Justin P. Gerke, and Leonid Kruglyak. 2015. “A Powerful New Quantitative Genetics Platform, Combining Caenorhabditis Elegans High-Throughput Fitness Assays with a Large Collection of Recombinant Strains.” G3 5 (5): 911–20.

Azzam, M. E., and I. D. Algranati. 1973. “Mechanism of Puromycin Action: Fate of Ribosomes after Release of Nascent Protein Chains from Polysomes.” Proceedings of the National Academy of Sciences of the United States of America 70 (12): 3866–69.

Balla, Keir M., Erik C. Andersen, Leonid Kruglyak, and Emily R. Troemel. 2015. “A Wild C. Elegans Strain Has Enhanced Epithelial Immunity to a Natural Microsporidian Parasite.” PLoS Pathogens 11 (2): e1004583.

Bates, Douglas, Martin Mächler, Ben Bolker, and Steve Walker. 2014. “Fitting Linear Mixed-Effects Models Using lme4.” arXiv [stat.CO]. arXiv. http://arxiv.org/abs/1406.5823.

Bendesky, Andres, and Cornelia I. Bargmann. 2011. “Genetic Contributions to Behavioural Diversity at the Gene-Environment Interface.” Nature Reviews. Genetics 12 (12): 809–20.

Bendesky, Andres, Jason Pitts, Matthew V. Rockman, William C. Chen, Man-Wah Tan, Leonid Kruglyak, and Cornelia I. Bargmann. 2012. “Long-Range Regulatory Polymorphisms Affecting a GABA Receptor Constitute a Quantitative Trait Locus (QTL) for Social Behavior in Caenorhabditis Elegans.” PLoS Genetics 8 (12): e1003157.

Bendesky, Andres, Makoto Tsunozaki, Matthew V. Rockman, Leonid Kruglyak, and Cornelia I. Bargmann. 2011. “Catecholamine Receptor Polymorphisms Affect Decision-Making in C. Elegans.” Nature 472 (7343): 313–18.

Bloom, Joshua S., Ian M. Ehrenreich, Wesley T. Loo, Thúy-Lan Võ Lite, and Leonid Kruglyak. 2013. “Finding the Sources of Missing Heritability in a Yeast Cross.” Nature 494 (7436): 234–37.

Borrello, Maria Grazia, Debora Degl’Innocenti, and Marco A. Pierotti. 2008. “Inflammation and Cancer: The Oncogene-Driven Connection.” Cancer Letters 267 (2): 262–70.

Boyd, Windy A., Marjolein V. Smith, and Jonathan H. Freedman. 2012. “Caenorhabditis Elegans as a Model in Developmental Toxicology.” Methods in Molecular Biology 889: 15–24.

Brady, Shannon C., Stefan Zdraljevic, Karol W. Bisaga, Robyn E. Tanny, Daniel E. Cook, Daehan Lee, Ye Wang, and Erik C. Andersen. 2019. “A Novel Gene Underlies Bleomycin-Response Variation in Caenorhabditis Elegans.” Genetics 212 (4): 1453–68.

Breitling, Rainer, Yang Li, Bruno M. Tesson, Jingyuan Fu, Chunlei Wu, Tim Wiltshire, Alice Gerrits, et al. 2008. “Genetical Genomics: Spotlight on QTL Hotspots.” PLoS Genetics 4 (10): e1000232.

Brem, Rachel B., Gaël Yvert, Rebecca Clinton, and Leonid Kruglyak. 2002. “Genetic Dissection of Transcriptional Regulation in Budding Yeast.” Science 296 (5568): 752–55.

Broman, Karl W., Hao Wu, Saunak Sen, and Gary A. Churchill. 2003. “R/qtl: QTL Mapping in Experimental Crosses.” Bioinformatics 19 (7): 889–90.

Brown, E. B., J. E. Layne, C. Zhu, A. G. Jegga, and S. M. Rollmann. 2013. “Genome-Wide Association Mapping of Natural Variation in Odour-Guided Behaviour in Drosophila.” Genes, Brain, and Behavior 12 (5): 503–15.

Chesmore, Kevin, Jacquelaine Bartlett, and Scott M. Williams. 2018. “The Ubiquity of Pleiotropy in Human Disease.” Human Genetics 137 (1): 39–44.

Cook, Daniel E., Stefan Zdraljevic, Joshua P. Roberts, and Erik C. Andersen. 2016. “CeNDR, the Caenorhabditis Elegans Natural Diversity Resource.” Nucleic Acids Research, October. https://doi.org/10.1093/nar/gkw893.

Cubillos, Francisco A., Eleonora Billi, Enikö Zörgö, Leopold Parts, Patrick Fargier, Stig Omholt, Anders Blomberg, Jonas Warringer, Edward J. Louis, and Gianni Liti. 2011. “Assessing the Complex Architecture of Polygenic Traits in Diverged Yeast Populations.” Molecular Ecology 20 (7): 1401–13.

Dasari, Shaloam, and Paul Bernard Tchounwou. 2014. “Cisplatin in Cancer Therapy: Molecular Mechanisms of Action.” European Journal of Pharmacology 740 (October): 364–78.

Doroszuk, Agnieszka, L. Basten Snoek, Emilie Fradin, Joost Riksen, and Jan Kammenga. 2009. “A Genome-Wide Library of CB4856/N2 Introgression Lines of Caenorhabditis Elegans.” Nucleic Acids Research 37 (16): e110.

Dorr, R. T. 1992. “Bleomycin Pharmacology: Mechanism of Action and Resistance, and Clinical Pharmacokinetics.” Seminars in Oncology 19 (2 Suppl 5): 3–8.

El-Assal, S. E. D., C. Alonso-Blanco, C. J. Hanhart, and M. Koornneef. 2004. “Pleiotropic Effects of the Arabidopsis Cryptochrome 2 Allelic Variation Underlie Fruit Trait-Related QTL.” Plant Biology 6 (4): 370–74.

Evans, Kathryn S., Shannon C. Brady, Joshua S. Bloom, Robyn E. Tanny, Daniel E. Cook, Sarah E. Giuliani, Stephen W. Hippleheuser, Mostafa Zamanian, and Erik C. Andersen. 2018. “Shared Genomic Regions Underlie Natural Variation in Diverse Toxin Responses.” Genetics, October. https://doi.org/10.1534/genetics.118.301311.

Fisher, R. A. n.d. The Genetical Theory of Natural Selection. Рипол Классик.

Fusari, Corina M., Rik Kooke, Martin A. Lauxmann, Maria Grazia Annunziata, Beatrice Enke, Melanie Hoehne, Nicole Krohn, et al. 2017. “Genome-Wide Association Mapping Reveals That Specific and Pleiotropic Regulatory Mechanisms Fine-Tune Central Metabolism and Growth in Arabidopsis.” The Plant Cell 29 (10): 2349–73.

García-González, Aurian P., Ashlyn D. Ritter, Shaleen Shrestha, Erik C. Andersen, L. Safak Yilmaz, and Albertha J. M. Walhout. 2017. “Bacterial Metabolism Affects the C. Elegans Response to Cancer Chemotherapeutics.” Cell 169 (3): 431–41.e8.

Ghosh, Rajarshi, Erik C. Andersen, Joshua A. Shapiro, Justin P. Gerke, and Leonid Kruglyak. 2012. “Natural Variation in a Chloride Channel Subunit Confers Avermectin Resistance in C. Elegans.” Science 335 (6068): 574–78.

Glater, Elizabeth E., Matthew V. Rockman, and Cornelia I. Bargmann. 2014. “Multigenic Natural Variation Underlies Caenorhabditis Elegans Olfactory Preference for the Bacterial Pathogen Serratia Marcescens.” G3 4 (2): 265–76.

Gratten, Jacob, and Peter M. Visscher. 2016. “Genetic Pleiotropy in Complex Traits and Diseases: Implications for Genomic Medicine.” Genome Medicine 8 (1): 78.

Gutteling, E. W., A. Doroszuk, J. A. G. Riksen, Z. Prokop, J. Reszka, and J. E. Kammenga. 2007. “Environmental Influence on the Genetic Correlations between Life-History Traits in Caenorhabditis Elegans.” Heredity 98 (4): 206–13.

Gutteling, E. W., J. A. G. Riksen, J. Bakker, and J. E. Kammenga. 2007. “Mapping Phenotypic Plasticity and Genotype-Environment Interactions Affecting Life-History Traits in Caenorhabditis Elegans.” Heredity 98 (1): 28–37.

Hasin-Brumshtein, Yehudit, Arshad H. Khan, Farhad Hormozdiari, Calvin Pan, Brian W. Parks, Vladislav A. Petyuk, Paul D. Piehowski, et al. 2016. “Hypothalamic Transcriptomes of 99 Mouse Strains Reveal Trans eQTL Hotspots, Splicing QTLs and Novel Non-Coding Genes.” eLife 5 (September). https://doi.org/10.7554/eLife.15614.

Hindorff, Lucia A., Praveen Sethupathy, Heather A. Junkins, Erin M. Ramos, Jayashri P. Mehta, Francis S. Collins, and Teri A. Manolio. 2009. “Potential Etiologic and Functional Implications of Genome-Wide Association Loci for Human Diseases and Traits.” Proceedings of the National Academy of Sciences of the United States of America 106 (23): 9362–67.

Huang, Yen-Tsung, Liming Liang, Miriam F. Moffatt, William O. C. M. Cookson, and Xihong Lin. 2015. “iGWAS: Integrative Genome-Wide Association Studies of Genetic and Genomic Data for Disease Susceptibility Using Mediation Analysis.” Genetic Epidemiology 39 (5): 347–56.

Jerison, Elizabeth R., Sergey Kryazhimskiy, James Kameron Mitchell, Joshua S. Bloom, Leonid Kruglyak, and Michael M. Desai. 2017. “Genetic Variation in Adaptability and Pleiotropy in Budding Yeast.” eLife 6 (August). https://doi.org/10.7554/eLife.27167.

Kammenga, Jan E., Agnieszka Doroszuk, Joost A. G. Riksen, Esther Hazendonk, Laurentiu Spiridon, Andrei-Jose Petrescu, Marcel Tijsterman, Ronald H. A. Plasterk, and Jaap Bakker. 2007. “A Caenorhabditis Elegans Wild Type Defies the Temperature-Size Rule Owing to a Single Nucleotide Polymorphism in Tra-3.” PLoS Genetics 3 (3): e34.

Kaplan, Ayse, Gulsen Akalin Ciftci, and Hatice Mehtap Kutlu. 2016. “Cytotoxic, Anti-Proliferative and Apoptotic Effects of Silver Nitrate against H-Ras Transformed 5RP7.” Cytotechnology 68 (5): 1727–35.

Kelley, Lawrence A., Stefans Mezulis, Christopher M. Yates, Mark N. Wass, and Michael J. E. Sternberg. 2015. “The Phyre2 Web Portal for Protein Modeling, Prediction and Analysis.” Nature Protocols 10 (6): 845–58.

Ketron, Adam C., William A. Denny, David E. Graves, and Neil Osheroff. 2012. “Amsacrine as a Topoisomerase II Poison: Importance of Drug-DNA Interactions.” Biochemistry 51 (8): 1730–39.

Keurentjes, Joost J. B., Jingyuan Fu, Inez R. Terpstra, Juan M. Garcia, Guido van den Ackerveken, L. Basten Snoek, Anton J. M. Peeters, Dick Vreugdenhil, Maarten Koornneef, and Ritsert C. Jansen. 2007. “Regulatory Network Construction in Arabidopsis by Using Genome-Wide Gene Expression Quantitative Trait Loci.” Proceedings of the National Academy of Sciences of the United States of America 104 (5): 1708–13.

Leamy, Larry J., Kari Elo, Merlyn K. Nielsen, Stephanie R. Thorn, William Valdar, and Daniel Pomp. 2014. “Quantitative Trait Loci for Energy Balance Traits in an Advanced Intercross Line Derived from Mice Divergently Selected for Heat Loss.” PeerJ 2 (May): e392.

Lee, Daehan, Heeseung Yang, Jun Kim, Shannon Brady, Stefan Zdraljevic, Mostafa Zamanian, Heekyeong Kim, et al. 2017. “The Genetic Basis of Natural Variation in a Phoretic Behavior.” Nature Communications 8 (1): 273.

Lin, Chi Hua Sarah, Jun Chen, Bruce Ziman, Shannon Marshall, Julien Maizel, and Michael S. Goligorsky. 2014. “Endostatin and Kidney Fibrosis in Aging: A Case for Antagonistic Pleiotropy?” American Journal of Physiology. Heart and Circulatory Physiology 306 (12): H1692–99.

Li, Yang, Olga Alda Alvarez, Evert W. Gutteling, Marcel Tijsterman, Jingyuan Fu, Joost A. G. Riksen, Esther Hazendonk, et al. 2006. “Mapping Determinants of Gene Expression Plasticity by Genetical Genomics in C. Elegans.” PLoS Genetics 2 (12): e222.

McGrath, Patrick T., Matthew V. Rockman, Manuel Zimmer, Heeun Jang, Evan Z. Macosko, Leonid Kruglyak, and Cornelia I. Bargmann. 2009. “Quantitative Mapping of a Digenic Behavioral Trait Implicates Globin Variation in C. Elegans Sensory Behaviors.” Neuron 61 (5): 692–99.

McGuigan, Katrina, Julie M. Collet, Elizabeth A. McGraw, Yixin H. Ye, Scott L. Allen, Stephen F. Chenoweth, and Mark W. Blows. 2014. “The Nature and Extent of Mutational Pleiotropy in Gene Expression of Male Drosophila Serrata.” Genetics 196 (3): 911–21.

McKay, J. K., J. H. Richards, and T. Mitchell-Olds. 2003. “Genetics of Drought Adaptation in Arabidopsis Thaliana: I. Pleiotropy Contributes to Genetic Correlations among Ecological Traits.” Molecular Ecology 12 (5): 1137–51.

Montecucco, Alessandra, Francesca Zanetta, and Giuseppe Biamonti. 2015. “Molecular Mechanisms of Etoposide.” EXCLI Journal 14 (January): 95–108.

Nikolova, Teodora, Wynand P. Roos, Oliver H. Krämer, Herwig M. Strik, and Bernd Kaina. 2017. “Chloroethylating Nitrosoureas in Cancer Therapy: DNA Damage, Repair and Cell Death Signaling.” Biochimica et Biophysica Acta 1868 (1): 29–39.

Orr, H. A. 2000. “Adaptation and the Cost of Complexity.” Evolution; International Journal of Organic Evolution 54 (1): 13–20.

Paaby, Annalise B., and Matthew V. Rockman. 2013. “The Many Faces of Pleiotropy.” Trends in Genetics: TIG 29 (2): 66–73.

Pavlides, Jennifer M. Whitehead, Zhihong Zhu, Jacob Gratten, Allan F. McRae, Naomi R. Wray, and Jian Yang. 2016. “Predicting Gene Targets from Integrative Analyses of Summary Data from GWAS and eQTL Studies for 28 Human Complex Traits.” Genome Medicine 8 (1): 84.

Peltier, Emilien, Anne Friedrich, Joseph Schacherer, and Philippe Marullo. 2019. “Quantitative Trait Nucleotides Impacting the Technological Performances of Industrial Saccharomyces Cerevisiae Strains.” Frontiers in Genetics 10 (July): 683.

Piperdi, Bilal, Yi-He Ling, Leonard Liebes, Franco Muggia, and Roman Perez-Soler. 2011. “Bortezomib: Understanding the Mechanism of Action.” Molecular Cancer Therapeutics.

R Core Team. 2017. “R: A Language and Environment for Statistical Computing.” Vienna, Austria: R Foundation for Statistical Computing. https://www.R-project.org/.

Reddy, Kirthi C., Erik C. Andersen, Leonid Kruglyak, and Dennis H. Kim. 2009. “A Polymorphism in Npr-1 Is a Behavioral Determinant of Pathogen Susceptibility in C. Elegans.” Science 323 (5912): 382–84.

Riedel, Christian G., Robert H. Dowen, Guinevere F. Lourenco, Natalia V. Kirienko, Thomas Heimbucher, Jason A. West, Sarah K. Bowman, et al. 2013. “DAF-16 Employs the Chromatin Remodeller SWI/SNF to Promote Stress Resistance and Longevity.” Nature Cell Biology 15 (5): 491–501.

Rockman, Matthew V., and Leonid Kruglyak. 2009. “Recombinational Landscape and Population Genomics of Caenorhabditis Elegans.” PLoS Genetics 5 (3): e1000419.

Rockman, Matthew V., Sonja S. Skrovanek, and Leonid Kruglyak. 2010. “Selection at Linked Sites Shapes Heritable Phenotypic Variation in C. Elegans.” Science 330 (6002): 372–76.

Rodriguez, Miriam, L. Basten Snoek, Joost A. G. Riksen, Roel P. Bevers, and Jan E. Kammenga. 2012. “Genetic Variation for Stress-Response Hormesis in C. Elegans Lifespan.” Experimental Gerontology 47 (8): 581–87.

Sasaki, Eriko, Florian Frommlet, and Magnus Nordborg. 2018. “GWAS with Heterogeneous Data: Estimating the Fraction of Phenotypic Variation Mediated by Gene Expression Data.” G3 8 (9): 3059–68.

Schmid, Tobias, L. Basten Snoek, Erika Fröhli, M. Leontien van der Bent, Jan Kammenga, and Alex Hajnal. 2015. “Systemic Regulation of RAS/MAPK Signaling by the Serotonin Metabolite 5-HIAA.” PLoS Genetics 11 (5): e1005236.

Seidel, Hannah S., Michael Ailion, Jialing Li, Alexander van Oudenaarden, Matthew V. Rockman, and Leonid Kruglyak. 2011. “A Novel Sperm-Delivered Toxin Causes Late-Stage Embryo Lethality and Transmission Ratio Distortion in C. Elegans.” PLoS Biology 9 (7): e1001115.

Seidel, Hannah S., Matthew V. Rockman, and Leonid Kruglyak. 2008. “Widespread Genetic Incompatibility in C. Elegans Maintained by Balancing Selection.” Science 319 (5863): 589–94.

Shimko, Tyler C., and Erik C. Andersen. 2014. “COPASutils: An R Package for Reading, Processing, and Visualizing Data from COPAS Large-Particle Flow Cytometers.” PloS One 9 (10): e111090.

Singh, Kapil Dev, Bernd Roschitzki, L. Basten Snoek, Jonas Grossmann, Xue Zheng, Mark Elvin, Polina Kamkina, et al. 2016. “Natural Genetic Variation Influences Protein Abundances in C. Elegans Developmental Signalling Pathways.” PloS One 11 (3): e0149418.

Sivakumaran, Shanya, Felix Agakov, Evropi Theodoratou, James G. Prendergast, Lina Zgaga, Teri Manolio, Igor Rudan, Paul McKeigue, James F. Wilson, and Harry Campbell. 2011. “Abundant Pleiotropy in Human Complex Diseases and Traits.” American Journal of Human Genetics 89 (5): 607–18.

Snoek, L. Basten, Helen E. Orbidans, Jana J. Stastna, Aafke Aartse, Miriam Rodriguez, Joost A. G. Riksen, Jan E. Kammenga, and Simon C. Harvey. 2014. “Widespread Genomic Incompatibilities in Caenorhabditis Elegans.” G3 4 (10): 1813–23.

Tingley, Dustin, Teppei Yamamoto, Kentaro Hirose, Luke Keele, and Kosuke Imai. 2014. “Mediation: R Package for Causal Mediation Analysis.” Journal of Statistical Software, Articles 59 (5): 1–38.

Tyler, Anna L., Dana C. Crawford, and Sarah A. Pendergrass. 2016. “The Detection and Characterization of Pleiotropy: Discovery, Progress, and Promise.” Briefings in Bioinformatics 17 (1): 13–22.

Viñuela, Ana, L. Basten Snoek, Joost A. G. Riksen, and Jan E. Kammenga. 2010. “Genome-Wide Gene Expression Regulation as a Function of Genotype and Age in C. Elegans.” Genome Research 20 (7): 929–37.

Wagner, Günter P., and Jianzhi Zhang. 2011. “The Pleiotropic Structure of the Genotype-Phenotype Map: The Evolvability of Complex Organisms.” Nature Reviews. Genetics 12 (3): 204–13.

White, Jacqueline K., Anna-Karin Gerdin, Natasha A. Karp, Ed Ryder, Marija Buljan, James N. Bussell, Jennifer Salisbury, et al. 2013. “Genome-Wide Generation and Systematic Phenotyping of Knockout Mice Reveals New Roles for Many Genes.” Cell 154 (2): 452–64.

Zamanian, Mostafa, Daniel E. Cook, Stefan Zdraljevic, Shannon C. Brady, Daehan Lee, Junho Lee, and Erik C. Andersen. 2018a. “Discovery of Genomic Intervals That Underlie Nematode Responses to Benzimidazoles.” PLoS Neglected Tropical Diseases 12 (3): e0006368.

Zamanian, Mostafa, Daniel E. Cook, Stefan Zdraljevic, Shannon C. Brady, Daehan Lee, Junho Lee, and Erik C. Andersen. 2018b. “Discovery of Genomic Intervals That Underlie Nematode Responses to Benzimidazoles.” PLoS Neglected Tropical Diseases 12 (3): e0006368.

Zdraljevic, Stefan, Bennett William Fox, Christine Strand, Oishika Panda, Francisco J. Tenjo, Shannon C. Brady, Tim A. Crombie, John G. Doench, Frank C. Schroeder, and Erik C. Andersen. 2019. “Natural Variation in C. Elegans Arsenic Toxicity Is Explained by Differences in Branched Chain Amino Acid Metabolism.” eLife 8 (April). https://doi.org/10.7554/eLife.40260.

Zdraljevic, Stefan, Christine Strand, Hannah S. Seidel, Daniel E. Cook, John G. Doench, and Erik C. Andersen. 2017. “Natural Variation in a Single Amino Acid Substitution Underlies Physiological Responses to Topoisomerase II Poisons.” PLoS Genetics 13 (7): e1006891.

Zhang, Lianqi, Lei Li, Liming Yan, Zhenhua Ming, Zhihui Jia, Zhiyong Lou, and Zihe Rao. 2018. “Structural and Biochemical Characterization of Endoribonuclease Nsp15 Encoded by Middle East Respiratory Syndrome Coronavirus.” Journal of Virology 92 (22). https://doi.org/10.1128/JVI.00893-18.

Zhao, Shanrong, Wai-Ping Fung-Leung, Anton Bittner, Karen Ngo, and Xuejun Liu. 2014. “Comparison of RNA-Seq and Microarray in Transcriptome Profiling of Activated T Cells.” PloS One 9 (1): e78644.

